# NSF/αSNAP2-mediated *cis*-SNARE complex disassembly precedes membrane fusion generating the cell plate during Arabidopsis cytokinesis

**DOI:** 10.1101/2022.04.29.490041

**Authors:** Misoon Park, Ulrike Mayer, Gerd Jürgens

## Abstract

Eukaryotic membrane fusion requires *trans*-SNARE complexes bridging the gap between adjacent membranes (Jahn and Scheller, 2006). Fusion between a transport vesicle and its target membrane transforms the *trans*-into a *cis*-SNARE complex. The latter interacts with the hexameric AAA^+^-ATPase NSF and its co-factor αSNAP, forming a 20S complex (Zhou et al., 2015; Zhao and Brunner, 2016). ATPase activity disassembles the SNARE complex into Qa-SNARE, which folds back onto itself, and its partners (Huang et al., 2019; Kim et al., 2021). Fusion of identical membranes has a different sequence of events (Baker and Hughson, 2016). The fusion partners each have *cis*-SNARE complexes to be broken up by NSF and αSNAP. The Qa-SNARE monomers are then stabilized by interaction with Sec1-type regulators (SM proteins) to form *trans*-SNARE complexes, as shown for the yeast vacuole (Baker et al., 2015). Membrane fusion in Arabidopsis cytokinesis is formally akin to vacuolar fusion (Müller and Jürgens, 2016). Membrane vesicles fuse with one another to form the partitioning membrane known as cell plate. C*is*-SNARE complexes of cytokinesis-specific Qa-SNARE KNOLLE and its SNARE partners are assembled at the ER and delivered by traffic via Golgi/TGN to the cell division plane (Karnahl and Park et al., 2017). SM protein KEULE is required for the formation of *trans*-SNARE complexes between adjacent membrane vesicles (Park et al., 2012). Here, we identify the missing NSF-type AAA^+^-ATPase and its adaptor αSNAP2 required for disassembly of KNOLLE *cis*-SNARE complexes. In addition, we show that NSF is also required for other trafficking pathways and interacts with the respective Q-SNAREs. In conclusion, the SNARE complex disassembly machinery is conserved in plants and plays a unique essential role in cytokinesis.

## Results and discussion

N-ethylmaleimide (NEM)-sensitive factor (NSF aka Sec18p; Novick et al., 1980) and its adaptor alpha-soluble NSF-associated protein 2 (αSNAP2 aka Sec17p; Novick et al., 1980) are both encoded by single-copy genes in Arabidopsis (AT4G04910 and AT3G56190, respectively). A putative αSNAP2 paralog, αSNAP1 (AT3G56450), is encoded in Arabidopsis. However, compared to αSNAP2, αSNAP1 is larger (44 kDa vs. 33 kDa) and is less similar to yeast and mammalian αSNAP proteins (Figure S1A), suggesting that αSNAP1 might be involved in other processes, and will not be considered here. NSF and/or αSNAP2 have been identified, by immunoprecipitation-mass spectrometry analysis, as potential interactors of several Qa-SNARE proteins on endomembranes but not of cytokinesis-specific Qa-SNARE KNOLLE nor plasma membrane-localized Qa-SNAREs (Fujiwara et al., 2014). In the presence of recombinant mammalian NSF and αSNAP proteins, however, some KNOLLE protein from detergent-solubilized microsomal fractions was recruited to 20S particles that cofractionated with NSF by glycerol gradient sedimentation (Rancour et al., 2002). The only NSF mutants described in Arabidopsis display mild leaf serration defects (Tang et al., 2020) or cause abnormal Golgi morphology (Tanabashi et al., 2018). By contrast, CRISPR/Cas9-generated αSNAP pre-mature stop codon mutants appear to be gametophytically lethal (Liu et al, 2021). There are no T-DNA insertion knockout alleles of NSF and CRISPR/Cas9 deletions of NSF appeared not to be transmissible (M.P and G.J, unpublished observation), suggesting that knockouts of αSNAP2-interacting NSF might also be gametophytically lethal (Liu et al, 2021). We therefore generated transgenic plants that overexpressed engineered dominant-negative variants conditionally. The variants NSF_E326Q_ and αSNAP2_L288A_ were designed by homology to known variants of the mammalian orthologs NSF_E329Q_ and αSNAP_L294A_, respectively, impairing ATPase activity and thus disassembly of the *cis*-SNARE complex, but not altering the overall protein structures (Figure 1A; Suppl. Figure S1B-E). Mammalian NSF_E329Q_ disables ATP hydrolysis by the D1 domain of NSF, which exerts a dominant-negative effect on all NSF subunits in the 20S complex (Whiteheart et al., 1994; Dalal et al., 2004). Bovine αSNAP_L294A_ still binds to NSF but inhibits its ATPase activity, preventing VAMP (aka R-SNARE) dissociation from the 20S complex (Barnard et al., 1997). We used the RPS5A-GAL4>>UAS two-component system (Weijers et al., 2003) for strong expression of either dominant-negative protein in Arabidopsis embryogenesis, which resulted in embryo lethality (Figure 1B-G; Suppl. Figure S1F-K; Suppl. Table S1). Similarly, estradiol-induced expression (Karnahl and Park et al., 2017) in germinating seeds caused seedling lethality, preventing any further development (Figure 1H-I; Suppl. Figure S1L-O). The comparable effects of the dominant-negative variants are consistent with the interaction of NSF and αSNAP2 in planta, as shown by co-immunoprecipitation from extracts of doubly transgenic plants (Figure 1J-K) and by colocalization of the two proteins in seedling roots (Suppl. Figure S2I-N). Thus, NSF and αSNAP2 play essential roles in Arabidopsis.

**Figure 1.**
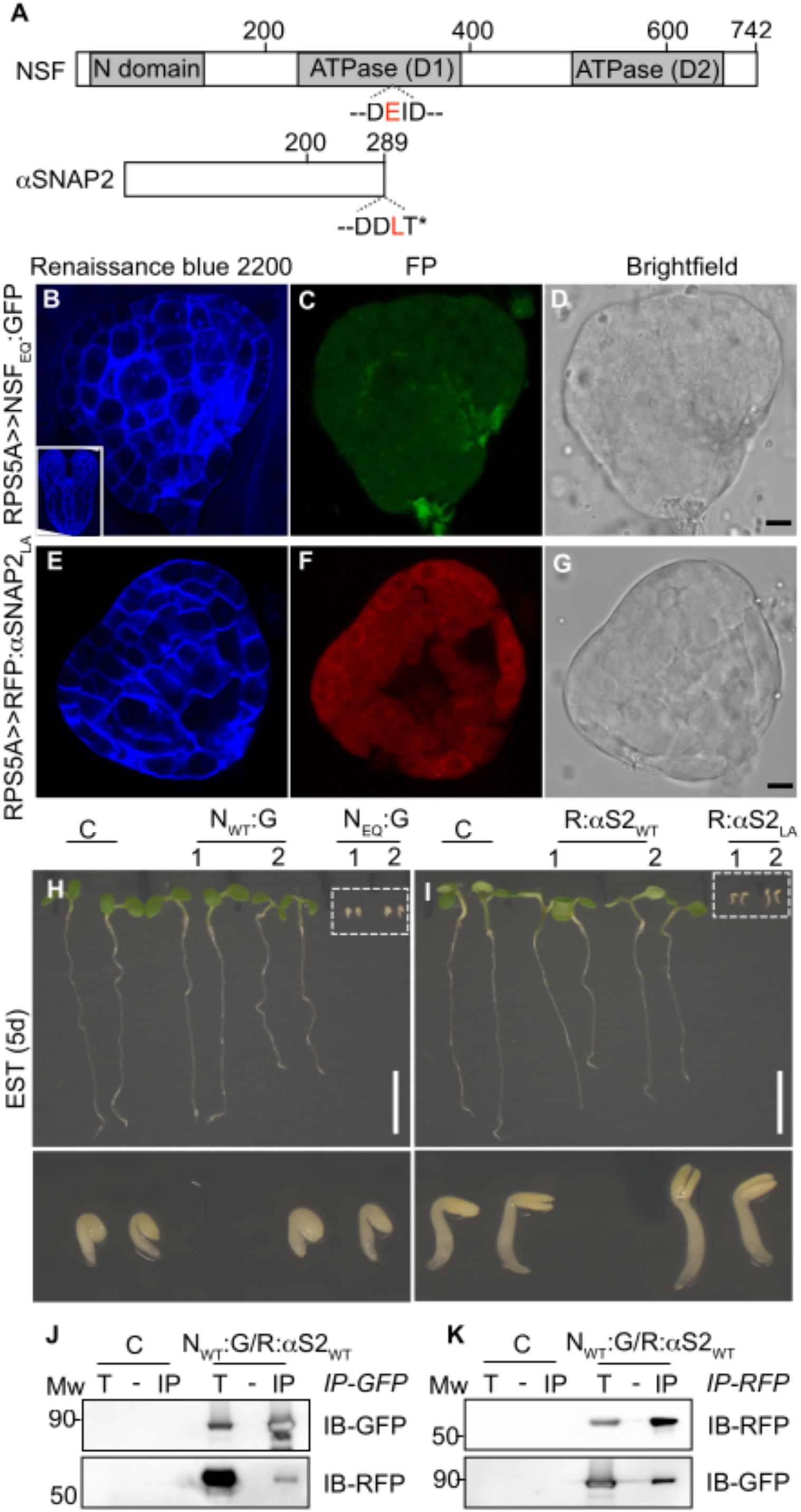
Interaction of NSF with αSNAP2 and impact of their dominant-negative variants on Arabidopsis development. (**A**) Schematic of NSF (*top*) and αSNAP2 (*bottom*). NSF domains: N domain for interaction with αSNAP2; ATPase (D1), (D2) in D1 and D2 regions. Numbers, amino acid positions. Residues altered in dominant-negative variants highlighted in red: E_326_ (NSF), L_288_ (αSNAP2). (**B-G**) Abnormal embryo phenotypes caused by two-component expression of dominant-negative NSF_E326Q_ (**B-D**) or αSNAP2_L288A_ (**E-G**). Inset (**B**), control embryo; (**B, E**) cell-wall staining with Renaissance blue 2200; (**C, F**) FP, fluorescent protein fusion (**C**, GFP; **F**, RFP) expression; (**D, G**) bright-field Images; scale bars, 10 μm. (**H-I**) Seedling phenotypes caused by expression of dominant-negative NSF_E326Q_ (**H**) or αSNAP2_L288A_ (**I**) induced by estradiol during seed germination and analyzed after 5 days. Scale bars, 1 cm. *Below*: Boxed areas at higher magnification. C, non-transformed wild-type; two transgenic lines (1, 2) each of N_WT_:G, NSF:GFP; N_EQ_:G, NSF_E326Q_:GFP; R:αS_WT_, RFP:αSNAP2; R:αS_LA_, RFP:αSNAP2_L288A_. See Figure S1K-L for control seedlings grown in DMSO. (**J-K**) Reciprocal co-immunoprecipitation between NSF:GFP and RFP:aSNAP2. T, total extract; IP, immunoprecipitate. IB, immunoblot with antibody (*right*); Mw, size markers (kDa, *left*). C, non-transformed wild-type extract; N_WT_:G/R:αS2_WT_, extract from transgenic line expressing GFP-tagged NSF and RFP-tagged αSNAP2 (wild-type forms); IP-GFP (**J**), IP-RFP (**K**), immunoprecipitation with anti-GFP or anti-RFP beads.

Subcellular localization analysis showed that wild type NSF was mainly located in the cytosol, at endosomes including the TGN and in the plane of cell division (Suppl. Figures S2A and S2C-E) whereas the dominant-negative NSF_E326Q_ protein was preferentially detected at membranes, and most strongly at the plasma membrane, rather than in the cytosol (Suppl. Figures S2B and S2F-H). This differential detection suggests that the dominant-negative NSF_E326Q_ variant might interact more strongly or longer than the wild-type form with some membrane proteins (compare Suppl. Figure S2B, F with S2A, C).

NSF and αSNAP2 are encoded by single-copy genes in Arabidopsis and might thus play a role in diverse trafficking pathways. We therefore examined whether the dominant-negative variant NSF_E326Q_ affected the subcellular distribution of trafficking markers. We used estradiol-inducible expression of both the trafficking marker and the possibly interfering dominant-negative variant NSF_E326Q_ in order to easily detect any change in the subcellular localization of the marker. The secretory marker secRFP accumulated in the extracellular space of seedling roots overexpressing the wild-type form of NSF (Figure 2A-C; Richter et al., 2014). By contrast, secRFP was retained inside the cell in seedling roots overexpressing the dominant-negative NSF_E326Q_ variant (Figure 2D-F). The accumulation of the marker underneath the plasma membrane and the preferential accumulation of the dominant-negative variant NSF_E326Q_ at the plasma membrane suggest that the dominant-negative NSF variant interferes with the fusion of the secretory vesicles with the plasma membrane. Similarly, the endocytic marker SynaptoRed™ C2, which is equivalent to FM4-64, was retained at the plasma membrane, rather than accumulating in endosomal dots (Suppl. Figure S3A-F). This effect was striking in seedling roots treated with the fungal toxin brefeldin A (BFA) which causes aggregation of endosomal membranes into large BFA compartments (Geldner et al., 2001). These SynaptoRed™ C2-positive BFA bodies were present in roots overexpressing the wild-type form of NSF but largely absent from the roots overexpressing the dominant-negative NSF_E326Q_ variant (Suppl. Figure S3G-L). The effect on endocytosis might be an indirect consequence of preventing vesicle fusion with the plasma membrane. Compared to these clear-cut effects, the consequences of NSF ATPase inhibition on vacuolar trafficking were less obvious. Soluble cargo AFVY:RFP (signal peptide:RFP:AFVY phaseolin vacuolar sorting sequence) was properly targeted to the vacuole (Richter et al., 2014), regardless of the overexpression of either the wild-type or the dominant-negative variant of NSF (Suppl. Fig. S3M-R).

**Figure 2.**
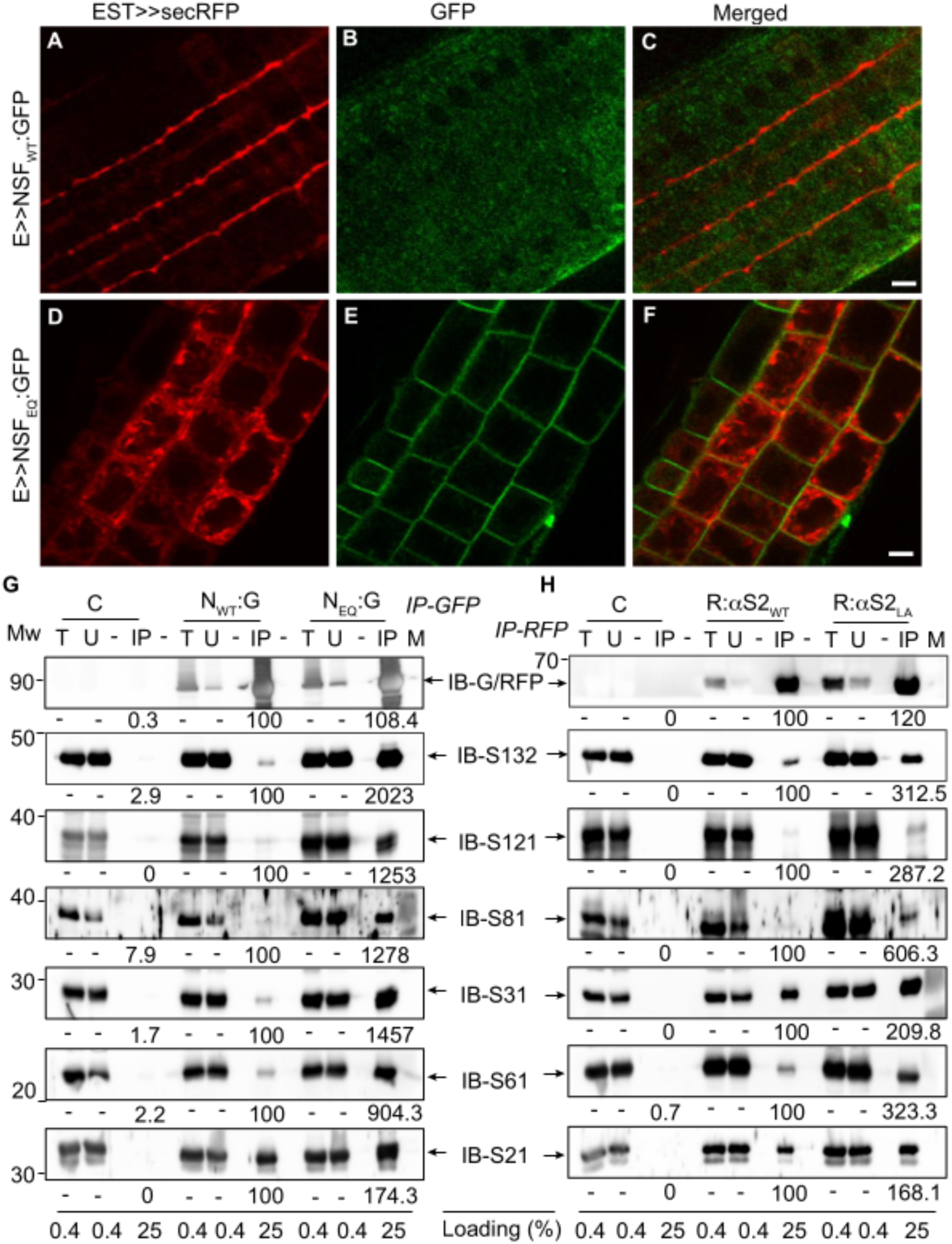
Interference of dominant-negative NSF_E326Q_ with secretory traffic and increased interaction between Q-SNAREs and the dominant -negative variants of NSF and αSNAP2. (**A-F**) Subcellular localization of secretory marker secRFP in root cells expressing the wild-type form of NSF (**A-C**) or the dominant-negative variant NSF_E326Q_ (**D-F**). (**A, D**) secRFP; (**B, E**) GFP-tagged NSF; (**C, F**) merged images; scale bars, 5 μm. (**G-H**) Co-immunoprecipitation of Q-SNAREs with (**G**) anti-GFP (IP-GFP) or (**H**) anti-RFP beads (IP-RFP). Extracts from non-transformed wild-type (C) or transgenic seedlings: NSF:GFP (N_wt_:G), NFS_E326Q_:GFP (N_EQ_:G), RFP:αSNAP2 (R:αS2_WT_), RFP:αSNAP2_L288A_ (R:αS2_LA_). Protein blots were probed with antibodies indicated on the right; sizes (Mw) on the left (kDa). Quantitative band intensity values are given in % relative to the value of the respective wild-type form of NSF or αSNAP2 (set at 100%). M, marker; IP, immunoprecipitate; T, total extract; U, unbound. IB, immunoblots with antibodies against Q-SNAREs: S132, SYP132 (mainly plasma membrane); S121, SYP121 (plasma membrane); S81, SYP81 (ER); S31, SYP31 (Golgi); S61, SYP61 (TGN); S21, SYP21 (MVB, vacuole).

Consistent with the deleterious effects on secretory and endocytic pathways, the dominant-negative variants NSF_E326Q_ and αSNAP_L188A_ displayed increased interaction, compared to wild-type forms of NSF and αSNAP2, with Q-SNAREs involved in diverse trafficking pathways, ranging - with one exception - from 9-fold to 20-fold and 2-fold to 12-fold, respectively (Figure 2G-H). Almost no increase was detected for SYP21 (aka as PEP12) which plays a role in vacuolar trafficking (Shirakawa et al., 2010; Uemura and Ueda, 2014). This is in line with the observation that blocking NSF ATPase activity leaves traffic to the vacuole seemingly undisturbed (see Suppl. Figure S3M-R).

In plant cytokinesis, membrane vesicles are delivered to the plane of cell division where they fuse with one another to form the partitioning cell plate (Samuels et al., 1995; Weizenegger et al., 2000). This fusion process requires the cytokinesis-specific Qa-SNARE KNOLLE which resides on the membrane vesicles and accumulates at the forming cell plate (Lukowitz et al., 1996; Lauber et al., 1997). To determine whether cytokinesis is compromised when ATPase activity of NSF is inhibited, we analyzed the subcellular localization of Qa-SNARE KNOLLE in embryos counterstained for nuclei (Figure 3). Embryos overexpressing the wild-type version of NSF displayed the normal arrangement of KNOLLE in dividing cells (Figure 3A-C). By contrast, embryos overexpressing dominant-negative NSF_E326Q_ showed wider bands of KNOLLE-positive material between daughter nuclei in dividing cells (Figure 3D-F). We detected a similar abnormality of KNOLLE distribution between the forming daughter nuclei in embryos expressing dominant-negative αSNAP2_L288A_ (Figure 3G-L). These results suggest that NSF activity is required for normal progression of cytokinesis.

**Figure 3.**
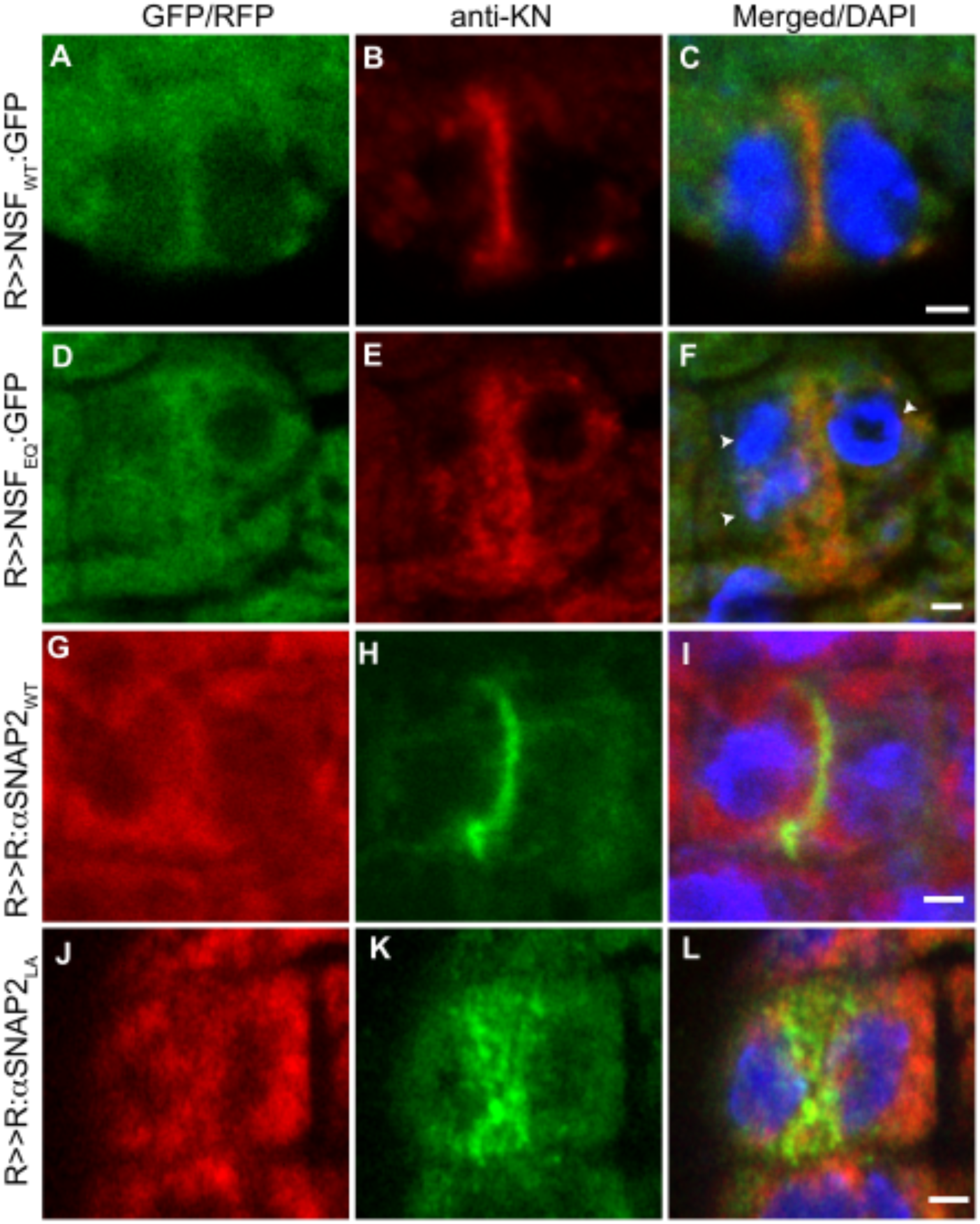
Cytokinesis defects caused by expression of dominant-negative NSF_E326Q_ or αSNAP_L288A_. Dividing cells from embryos expressing fluorescent (**A-C**) NSF, (**D-F**) NSF_E326Q_, (**G-I**) αSNAP2, (**J-L**) αSNAP2_L288A_ with the RPS5A:GAL4>>UAS system. (**A, D**) GFP; (**G, J**) RFP; (**B, E, H, K**) KNOLLE detected with antiserum; (**C, F, I, L**) merged images and nuclei stained with DAPI; scale bars, 5 μm. Arrowheads (**F**), 3 nuclei in dividing cell.

Cytokinesis-specific Qa-SNARE KNOLLE forms two *cis*-SNARE complexes at the ER, one involving Qbc-SNARE SNAP33 and R-SNARE VAMP721 or VAMP722 whereas the other complex comprises Qb-SNARE NPSN11, Qc-SNARE SYP71 and the same R-SNARE (El Kasmi and Krause et al., 2013; Karnahl and Park et al., 2017). These KNOLLE cis-SNARE complexes are trafficked from the ER via Golgi/TGN to the plane of cell division (Karnahl and Park et al., 2017). We probed by co-immunoprecipitation the interaction of NSF and αSNAP2 with these two complexes (Figure 4A-B). Compared to the wild-type form, the dominant-negative NSF_E326Q_ variant co-precipitated KNOLLE 38-fold and all its SNARE partners in the two complexes 18-fold to 51-fold more effectively (Figure 4A). Similarly, dominant-negative αSNAP_L288A_ also interacted up to 6-fold more strongly than its wild-type form with KNOLLE and all its SNARE partners (Figure 4B). These results provide a molecular explanation for the deleterious effects that the inhibition of the NSF ATPase has on cytokinesis. In this context, it should be noted that the SNARE partners of KNOLLE are shared with plasma membrane-localized Qa-SNAREs SYP132 and SYP121 (aka PEN1) which also interacted strongly with the dominant-negative variants NSF_E326Q_ and αSNAP2_L288A_ (see Figure 2G-H; Park et al., 2018; Kwon et al., 2008). This strongly increased interaction might explain the prominent localization of the dominant-negative NSF_E326Q_ variant at the membranes including cell division plane and plasma membrane. Thus, Arabidopsis NSF ATPase and its adaptor αSNAP2 appear to play essential roles in *cis*-SNARE complex disassembly in cell-plate formation and other membrane fusion processes.

**Figure 4.**
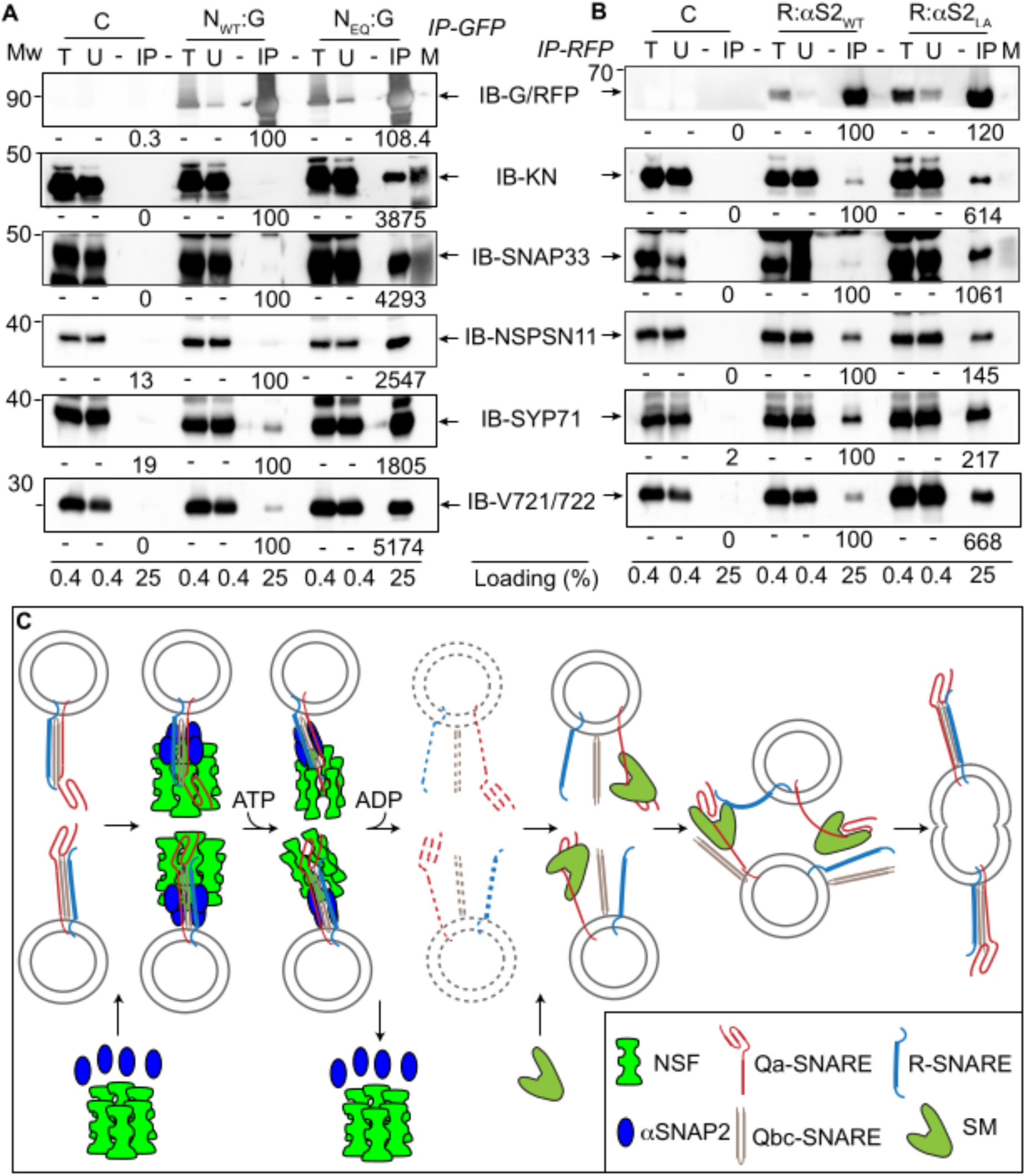
Increased interaction of dominant-negative variants of NSF and αSNAP2 with cytokinesis-specific Qa-SNARE KNOLLE and its SNARE partners, and role of NSF and αSNAP2 in cell-plate formation. (**A-B**) Co-immunoprecipitation of Qa-SNARE KNOLLE and its SNARE partners with (**A**) anti-GFP (IP-GFP) or (**B**) anti-RFP beads (IP-RFP). Extracts from non-transformed wild-type (C) or transgenic seedlings: NSF:GFP (N_wt_:G), NFS_E326Q_:GFP (N_EQ_:G), RFP:αSNAP2 (R:αS2_WT_), RFP:αSNAP2_L288A_ (R:αS2_LA_). Protein blots were probed with antibodies indicated on the right; sizes (Mw) on the left (kDa). Quantitative band intensity values are given in % relative to the value of the respective wild-type form of NSF or αSNAP2 (set at 100%). M, marker; IP, immunoprecipitate; T, total extract; U, unbound. IB, immunoblots with antibodies against SNAREs: KN, KNOLLE; SNAP33 (Qbc-SNARE); NPSN11 (Qb-SNARE), SYP71 (Qc-SNARE); V721/722, VAMP721/VAMP722 (R-SNARE). Note that the immunoblot images of anti-GFP (**A**) and anti-RFP (**B**) were the same as in Figure 2G and 2H, respectively, as the experiment was done at the same time. (**C**) A mechanistic model of membrane fusion in Arabidopsis cell plate formation. TGN-derived identical vesicles are delivered to the forming cell plate along the microtubules of the phragmoplast (not drawn). Hexameric NSF (bright green) and four copies of αSNAP2 (bright blue) form a 20S particle with the *cis*-SNARE complex (KNOLLE, Qa-SNARE, red; SNAP33, Qbc-SNARE, sand brown; VAMP721 or 722, R-SNARE, dark blue) on each vesicle. By ATP hydrolysis through NSF AAA^+^-ATPase activity, the *cis*-SNARE complex is dissembled into the monomeric SNAREs (broken outlines, indicating a very transient state). SM protein KEULE (pale green) interacts with a monomeric fusion-competent KNOLLE and accelerates *trans*-SNARE complex formation with KNOLLE-interacting SNARE partners residing on adjacent vesicles, resulting in vesicles fusion.

Plant cytokinesis is unique among the eukaryotes in that the non-plant plasma-membrane constriction driven by a contractile actomyosin ring and the subsequent ESCRT-mediated abscission of the intercellular bridge are replaced by the targeted delivery of specific membrane vesicles and their subsequent fusion to generate the partitioning membrane de novo (Vietri et al., 2020; Frémont and Echard, 2018; Müller and Jürgens, 2016). Our current study supports a mechanistic model of how membrane fusion is regulated in plant cytokinesis (Figure 4C). Hexameric NSF ATPase associates via 4 copies of its adaptor αSNAP2 (adapted from Chang et al., 2012 and Zhou et al., 2015) with KNOLLE *cis*-SNARE complexes on Golgi/TGN-derived membrane vesicles accumulating in the plane of cell division. ATPase activity disassembles the KNOLLE complex into monomeric SNAREs. The SM protein KEULE, which is also present in the plane of cell division and required for cell-plate formation, then captures monomeric KNOLLE, thus preventing its transition to the inactive closed conformation (Park et al., 2012). Instead, KEULE might form a template complex, bringing the N-terminal regions of the SNARE domains of Qa-SNARE KNOLLE and R-SNARE VAMP721 into close proximity while keeping their C-terminal regions separated. This kind of three-protein interaction is inferred from the single-molecule force microscopy analysis of the interaction of SM protein Munc18-1 with neuronal syntaxin and VAMP2 (Jiao et al., 2018). The same template mechanism has also been demonstrated for Munc18-3 and vacuolar SM protein Vps33 with their cognate SNARE partners and thus appears to be conserved among SM proteins (Jiao et al., 2018). It is therefore tempting to speculate that an equivalent templating complex would initiate the formation of *trans*-SNARE complexes required for the fusion of cytokinetic vesicles with one another to form the cell plate.

## Materials and methods

### Plants lines and growth conditions

Plants on soil or on plant medium (0.1 % MES, 1% sucrose, 2.15 g/l Murashige and Skoog, pH5.6) were grown in long days (16hr/8hr) at 23°C. Transgenic plants were generated by transforming wild type *Arabidopsis thaliana* (Col-0) with the *Agrobacterium tumefaciens*-mediated flower-dipping method (Clough and Bent, 1998) using. T1 plants bearing pMDC7-driven (Karnarhl and Park et al., 2017) or pGIIB-UAS driven transgenes (Park et al, 2012) were selected with 2 μg/ml hygromycin B (Duchefa) or 15 μg/ml PPT (phosphinothricin, Sigma-Aldrich), respectively. Plants bearing pMDC7::NSF_WT_ or pMDC7::NSF_E326Q_:GFP were crossed with plants bearing pMDC7::mRFP:αSNAP2_WT_, a1-RFP (Dettmer et al., 2006), pMDC7::secRFP (Richter et al., 2014) or pMDC7::AFVY:RFP (Richter et al., 2014). The resulting F1 was analyzed. Plants bearing pGIIB-UAS::NSF_WT_, pGIIB-UAS::NSF_E326Q_:GFP, pGIIB-UAS::mRFP:αSNAP2_WT_ or pGIIB-UAS::mRFP:αSNAP2_L288A_ were crossed with plants of the activator line pGIIKan-RPS5A::GAL4 (Weijers et al., 2003) for the analysis of the resulting F1.

### Molecular cloning

The coding sequences corresponding to *NSF*_*E326Q*_ and *αSNAP2*_*L288A*_ were commercially synthesized (BaseClear, Leiden, NL). *NSF*_*E326Q*_*:GFP* and *mRFP:αSNAP2*_*L288A*_ were generated by combining PCR-amplified *GFP* or *mRFP* together with PCR-amplified *NSF*_*E326Q*_ or *αSNAP2*_*L288A*_, respectively, using the primers listed in Table S2. *NSF*_*WT*_*:GFP* and *mRFP:αSNAP2*_*WT*_ were generated by PCR, using *NSF*_*E326Q*_*:GFP* and *mRFP:αSNAP2*_*L288A*_ as templates with the rescuing primers listed in Table S2. These PCR products were subsequently subcloned into *pDONOR221* and *pMDC7* gateway plasmids. For cloning downstream of the *UAS* element, *NSF*_*WT*_: *GFP* or *NSF*_*E326Q*_*:GFP* and *mRFP:αSNAP2*_*WT*_ or *mRFP:αSNAP2*_*L288A*_ were PCR-amplified with the primers listed in Table S2, digested with *Xho*I and *Hin*dIII or *Eco*RV and subcloned into the *pGIIB-UAS-tNOS* plasmid. See the Table S2 for the primers sequences.

### Chemical treatment

Five-day-old seedlings were transferred to media supplemented with 20 μM α-estradiol (EST, 20 mM stock in DMSO, Sigma-Aldrich) and incubated in a plant growth room with gentle shaking for 24 hours. 1 μM SynaptoRed™ C2 (1 mM stock in DMSO, Sigma-Aldrich) or 50 μM BFA (50 mM stock in DMSO/Ethanol, Invitrogen) was applied 30 minutes before the microscopic observation.

### Immunohistochemistry

Embryos or seedlings were fixed with 4% paraformaldehyde in MTSB (50 mM Pipes, 5 mM EGTA, 5 mM MgSO4, pH6.9∼7.0) solution for 1 hour, mounted on adhesive glass slides (manually prepared gelatin-coated slides and SUPERFROST^®^ PLUS (Thermo Scientific, USA), respectively, dried and stored at −20°C until used. Primary antiserum anti-KN (1:2000, rabbit) (Lauber et al., 1997) and secondary antibodies anti-rabbit Alex488 (1:600, Invitrogen), anti-rabbit Cy3 (1:600, Dianova) were applied. For staining the nucleus, 1 μg/ml DAPI (1mg/ml stock in H_2_O) was used. Renaissance blue 2200 staining was performed as described (Musielak et al., 2015). Fluorescent images were acquired with a confocal laser scanning microscope (SP8, Leica).

### Immunoprecipitation and immunodetection

Frozen seedlings were ground in liquid nitrogen (N_2_) and suspended in a chilled IP buffer (50 mM Tris pH7.5,150 mM NaCl, 1mM EDTA, 0.5% Triton X-100) supplemented with EDTA-free proteases cocktail (Roche diagnostics). Precleared lysates were incubated with GFP-trap or RFP-trap agarose beads (Chromotek, Munich, Germany) for 2 hours at 4°C with mild rotation. Five-times washed beads were suspended with 2x protein sample buffer, boiled for 5 minutes at 95°C and subjected to immunodetection. The membrane was developed with a chemiluminescence detection system (Fusion Fx7 Imager, PEQlab, Erlangen, Germany), using the detection solution (BM detection system, Roche diagnostics). The following antibodies were used: anti-GFP (1:1000-1:4000, mouse, Roche diagnostics), anti-RFP (1:1000, rat, Chromotek), anti-KN (1:4000, rabbit) (Lauber et al., 1997), anti-SNAP33 (1:5000, rabbit) (Heese et al., 2001), anti-NPSN11 (1:1500, rabbit) (Zheng et al., 2002), anti-SYP71 (1:2000, rabbit) (Sanderfoot et al., 2001), anti-VAMP721/722 (1:4000, rabbit) (Kwon et al., 2008), anti-SYP132 (1:4000, rabbit) (Park et al., 2018), anti-SYP121 (1:4000, rabbit) (Zhang et al., 2007), anti-SYP61 (1:1000, rabbit) (Sanderfoot et al., 2001), anti-SYP21 (1:1000, rabbit) (Sanderfoot et al., 1999), anti-SYP31 (1:1000, rabbit) (Rancour et al., 2002), anti-SYP81 (1:500, rabbit) (Bubeck et al., 2008), goat anti-mouse IgG-POD polyclonal antibody (1:4000, Sigma-aldrich), goat anti-rat IgG-POD polyclonal antibody (1:4000, Sigma-aldrich), goat anti-rabbit IgG-POD polyclonal antibody (1:20000, Agrisera).

### Protein structure prediction, intensity quantification and sequence alignment

The structure of NSF_E326Q_ was predicted using Arabidopsis NSF (Q9M0Y8) as template in SWISS-MODEL (http://swissmodel.expasy.org). The proteins structure of αSNAP2_WT_ and αSNAP2_L288A_ were predicted with the AlphaFold program (http://colab.research.google.come/github/deepmind/alphafold/blob/main/notebooks/AlphaFold.ipynb). Structures were superimposed in the PyMOL program (v0.99, Schrödinger, Inc.). For intensity quantification, the immunodetection images were quantified with the ImageJ Fiji program (NIH). For sequence alignment, the CLC Main Workbench (v20.0.1) program was used.

## Supplemental information

Supplemental Figure S1 (related to Figure 1).

Supplemental Figure S2 (related to Figure 1).

Supplemental Figure S3 (related to Figure 2).

Supplemental Table S1. Effects of dominant-negative NSF and αSNAP2 on seed viability.

Supplemental Table S2. List of primers used for cloning.

## Acknowledgments

We are indebted to Katerina Romanova for initial mutants screening, Sebastian Bednarek (UW, Madison) for anti-SYP31 antiserum, Karin Schumacher (Heidelberg University) for anti-SYP81 antiserum, Tomohiro Uemura (Ochanomizu University) and Akihiko Nakano (Riken center) for sharing T-DNA lines, and Martin Bayer and Farid El Kasmi for critical reading. This work is funded by the Deutsche Forschungsgemeinschaft (DFG JU 179/24-1 to. J.G.).

## Author contributions

Conceptualization, M.P. and G.J.; methodology, M.P. and U.M.; investigation, M.P. and U.M.; writing – original draft, M.P. and G.J.; writing – review & editing, M.P., U.M. and G.J.; funding acquisition, G.J.; resources, G.J.; supervision, G.J.

