## Supplemental Figures S1-S3, Supplemental Tables S1-S2 for "NSF/αSNAP2-mediated *cis*-SNARE complex disassembly precedes membrane fusion generating the cell plate during Arabidopsis cytokinesis"

**Inventory of supplemental information**

Supplemental Figure S1 (related to Figure 1).

Supplemental Figure S2 (related to Figure 1).

Supplemental Figure S3 (related to Figure 2).

Supplemental Table S1. Effects of dominant-negative NSF and αSNAP2 on seed viability.

Supplemental Table S2. List of primers used for cloning.

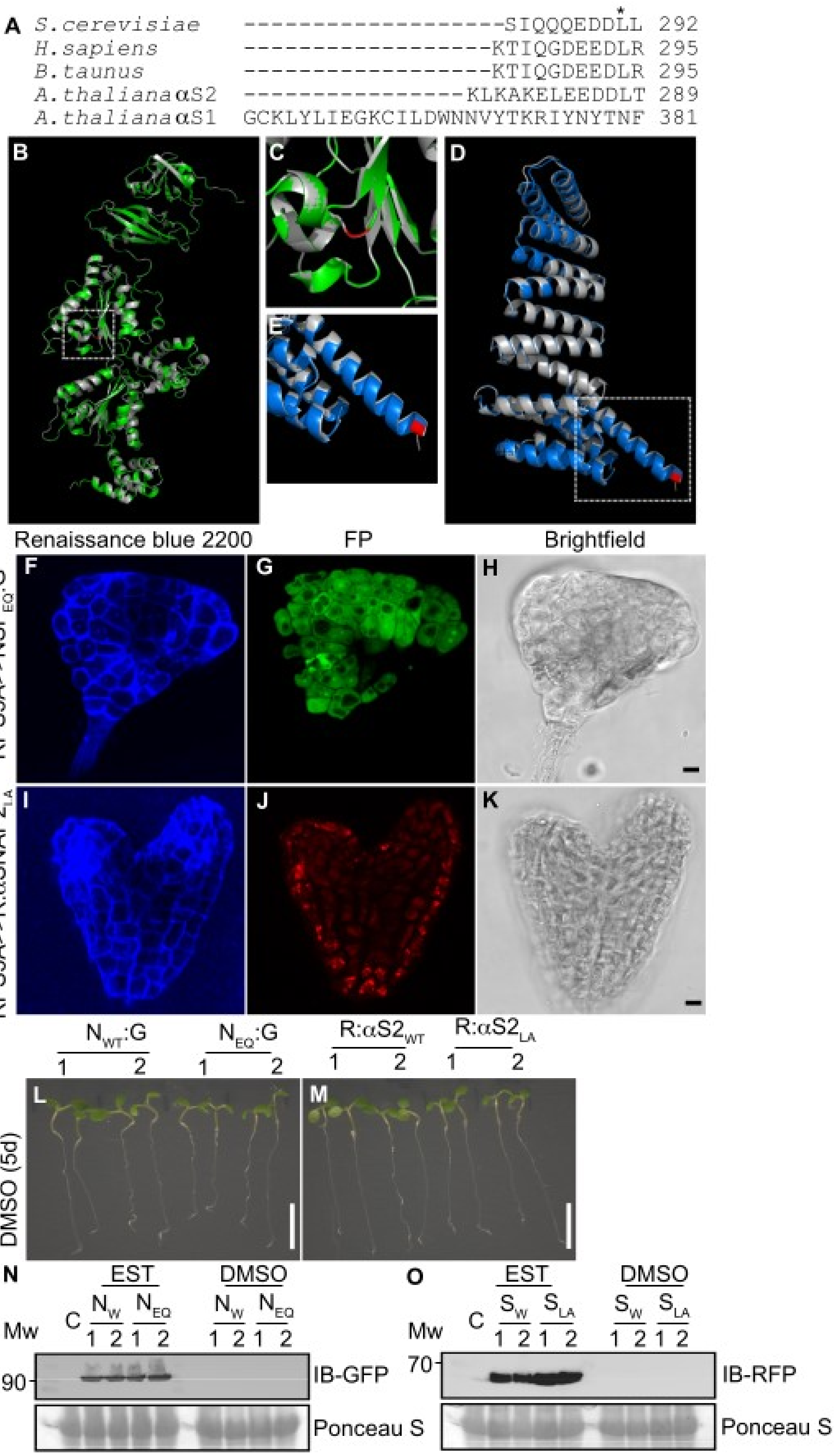
A S.cerevisiae --------------------SIQQQEDDLL 292
H.sapiens -------------------KTIQGDEEDLR 295
B.taunus -------------------KTIQGDEEDLR 295
A.thalianaαS2 -----------------KLKAKELEEDDLT 289
A.thalianaαS1 GCKLYLIEGKCILDWNNVYTKRIYNYTNF 381
B
C
D
E
Renaissance blue 2200
FP
Brightfield
F
G
H
RPS5A>>NSF$_{EQ}$:G
I
J
K
RPS5A>>R:αSNAP2$_{LA}$
N$_{WT}$:G
N$_{EQ}$:G
R:αS2$_{WT}$
R:αS2$_{LA}$
1 2 1 2 1 2 1 2
L
M
DMSO (5d)
N
EST
DMSO
C
N$_{W}$ N$_{EQ}$ N$_{W}$ N$_{EQ}$
Mw
1 2 1 2 1 2 1 2
90
IB-GFP
Ponceau S
O
EST
DMSO
C
S$_{W}$ S$_{LA}$ S$_{W}$ S$_{LA}$
Mw
1 2 1 2 1 2 1 2
70
IB-RFP
Ponceau S

**Figure S1 (related to Figure 1). Sequence alignment, predicted protein structures, embryo phenotypes, seedling controls and protein blots showing estradiol-induced expression**

**(A)** Alignment of C-terminal sequences of αSNAP-related proteins. *Arabidopsis thaliana* αSNAP1 (AT3G56450) and αSNAP2 (AT3G56190) were aligned with αSNAP from human (*Homo sapiens*, GenBank NP_003818), cattle (*Bos taurus*, GenBank AAB25812) and yeast (*Saccharomyces cerevisiae*, GenBank NP_009503) in the CLC Main Workbench program. Note that αSNAP1 is larger than the other αSNAPs and less similar to bovine αSNAP (E value 1e-29 vs. 2e-71 for αSNAP2). Asterisk marks the conserved leucine residue that was mutated to alanine in this study. Numbers indicate protein lengths (amino acid residues). (**B-E**) Structural models of NSF and αSNAP2. Predicted structures of wild type NSF (grey, **B** and **C**) and αSNAP2 (grey, **D** and **E**) were superimposed with those of $NSF_{E326Q}$ (green, **B** and **C**) and $αSNAP2_{L288A}$ (blue, **D** and **E**), respectively. The substituted residues of $NSF_{E326Q}$ and of $αSNAP2_{L288A}$ were colored in red (**B**-**E**). (**C** and **E**) Boxed areas in (**B** and **D**) at higher magnification. (**F-K**) Overview of phenotypes of late-stage degenerated embryos expressing RPS5A>>$NSF_{E326Q}$:GFP (**F-H**) or RPS5A>>RFP:$αSNAP2_{L288A}$ (**I-K**). (**F, I**) Cell-wall staining with Renaissance blue 2200; (**G, J**) FP, fluorescent protein fusion (**G**, GFP; **J**, RFP) expression; (**H, K**) bright-field Images; scale bars, 10 µm. (**L-M**) Seedlings grown in DMSO. Two transgenic lines (1, 2) each of $N_{WT}$:G, NSF:GFP; $N_{EQ}$:G, $NSF_{E326Q}$:GFP; R:$αS_{WT}$, RFP:αSNAP2; R:$αS_{LA}$, RFP:$αSNAP2_{L288A}$ were germinated and grown in DMSO-containing media for 5 days; scale bars, 1 cm. (**N-O**) Expression of transgene-encoded proteins. Proteins extracted from two transgenic seedlings lines (1, 2) incubated with EST (inducer) or DMSO (control) for 24 hours were immunoblotted (IB) with anti-GFP (**N**) and anti-RFP (**O**) antibodies. C, non-transformed wild type (Col-O); Nw, NSF:GFP; $N_{EQ,}$ $NSF_{E326Q}$:GFP;$S_{WT}$, RFP:αSNAP2; $S_{LA}$, RFP:$αSNAP2_{L288A.}$ Mw, size marker (kDa); Ponceau S, Ponceau S-stained membrane as loading control.

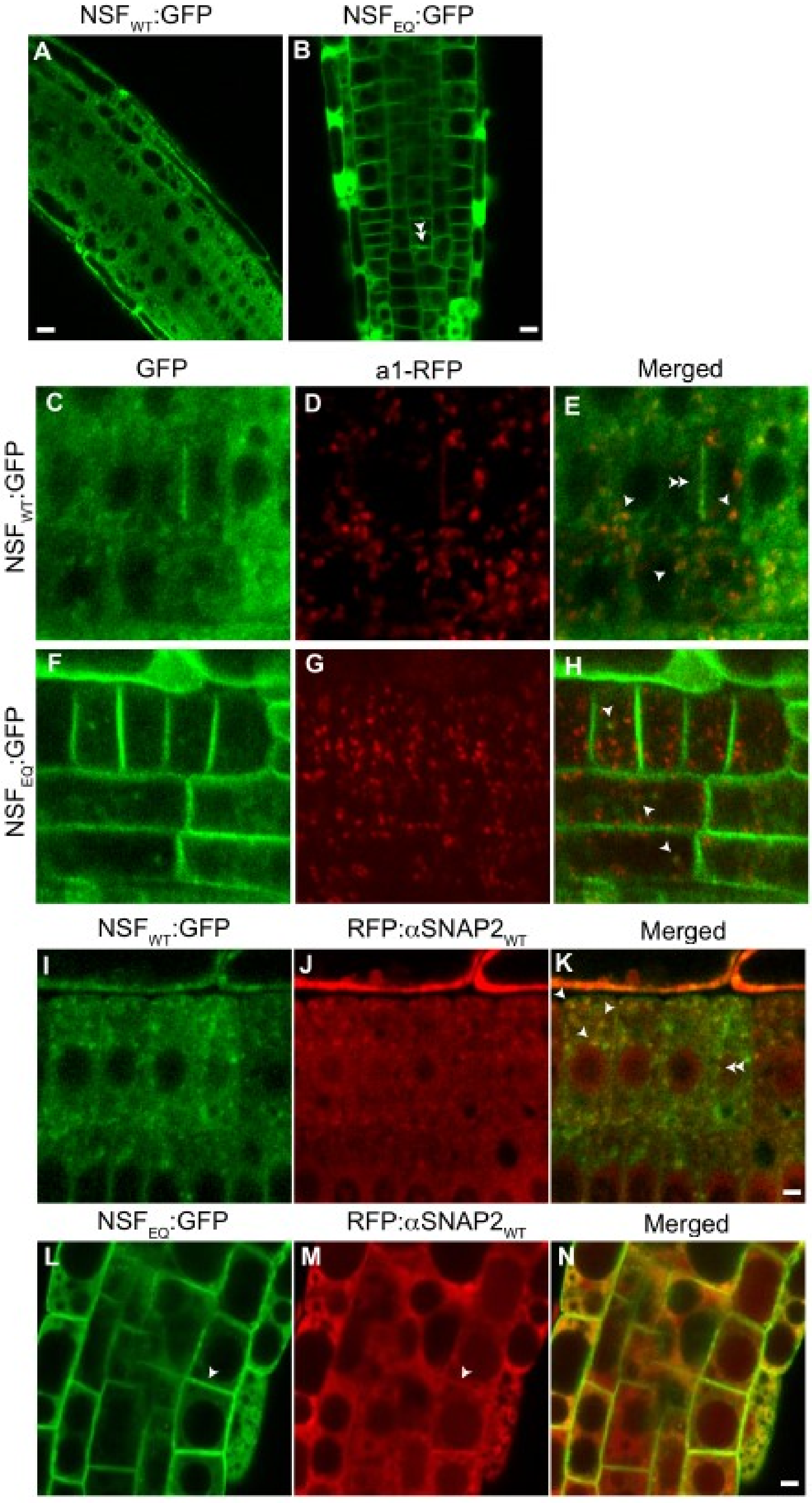
NSF$_{WT}$:GFP
NSF$_{EQ}$:GFP
A
B
GFP
a1-RFP
Merged
C
D
E
NSF$_{WT}$:GFP
F
G
H
NSF$_{EQ}$:GFP
NSF$_{WT}$:GFP
RFP:αSNAP2$_{WT}$
Merged
I
J
K
NSF$_{EQ}$:GFP
RFP:αSNAP2$_{WT}$
Merged
L
M
N

**Supplemental Figure S2 (related to Figure 1). Subcellular localization of NSF and αSNAP2**

(**A-B**) Live imaging of $NSF_{WT}$:GFP (**A**) and $NSF_{EQ}$:GFP (**B**) in seedling roots after 24 hours of EST induction. (**C-H**) Localization of NSF at endosomes. Live imaging of $NSF_{WT}$:GFP (**C**-**E**) and $NSF_{EQ}$:GFP (**F-H**) against TGN marker a1-RFP in seedling roots. Arrowheads indicate merged puncta signals of $NSF_{WT}$:GFP (**E**) or $NSF_{EQ}$:GFP (**H**) with al-RFP. (**I-N**) Live imaging of $NSF_{WT}$:GFP (**I-K**) or $NSF_{EQ}$:GFP (**L-N**) and RFP:$\alpha SNAP2_{WT}$ in seedling roots. Arrowheads indicate merged signals of $NSF_{WT}$:GFP (**K**) or $NSF_{EQ}$:GFP (**L** and **M**) with RFP:$aSNAP2_{WT}$. Double arrowheads indicate the cell division plane positively labeled with $NSF_{WT}$:GFP (**E** and **K**) or $NSF_{EQ}$:GFP (**B**). Scale bars, 5 µm (**A**, **B**, **E**, **H**, **K**, **N**).

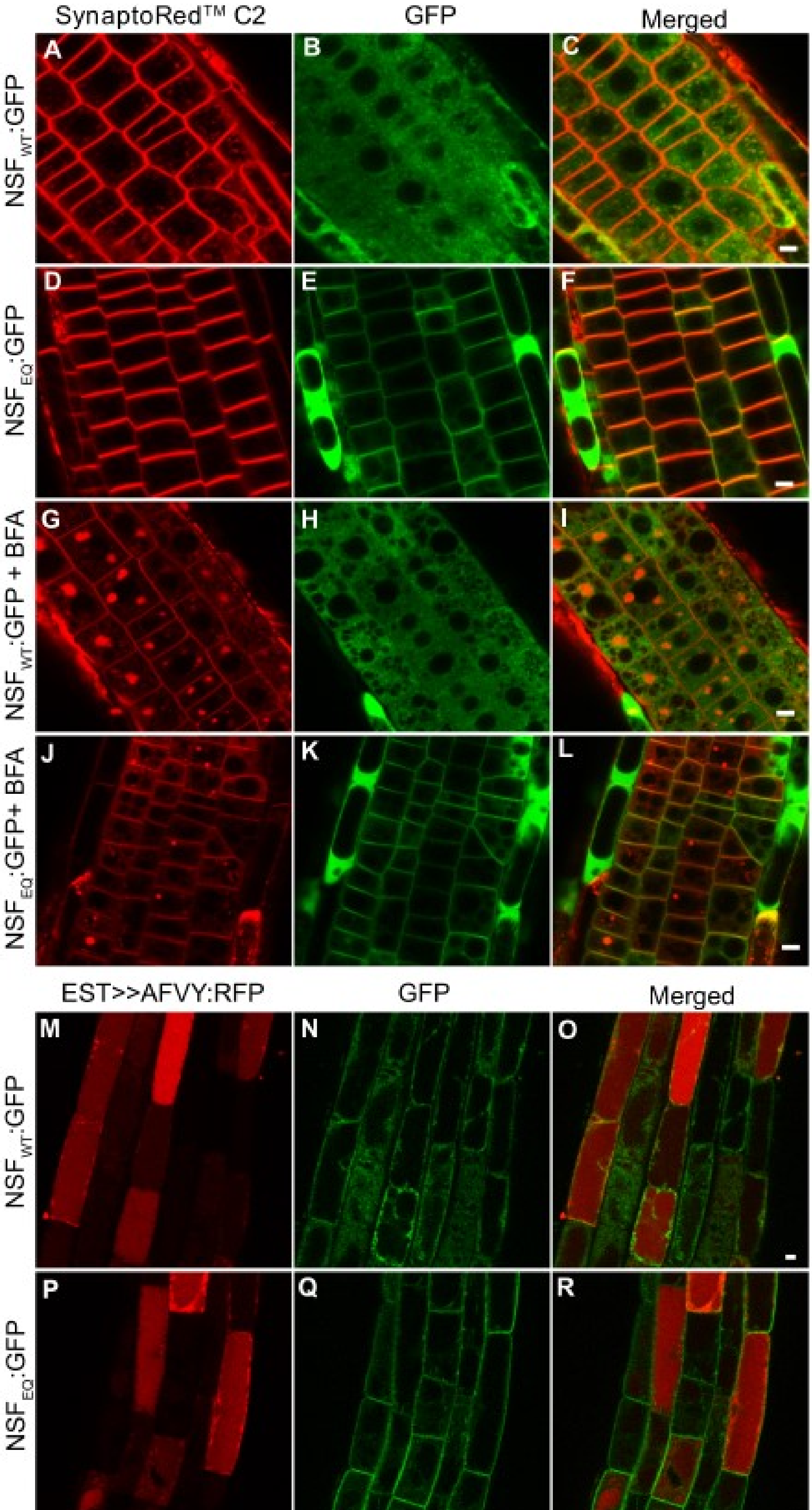
SynaptoRed™ C2
GFP
Merged
NSF$_{WT}$:GFP
NSF$_{EQ}$:GFP
NSF$_{WT}$:GFP + BFA
NSF$_{EQ}$:GFP+ BFA
A
B
C
D
E
F
G
H
I
J
K
L
EST>>AFVY:RFP
GFP
Merged
M
N
O
NSF$_{WT}$:GFP
P
Q
R
NSF$_{EQ}$:GFP

**Supplemental Figure S3 (related to Figure 2). Effect of dominant-negative NSF on trafficking markers.**

(**A-L**) Effect on endocytosis. (**A-F**) Live imaging of $NSF_{WT}$:GFP (**A-C**) or $NSF_{EQ}$:GFP (**D-F**) and endocytic tracer SynaptoRed™ C2 in seedling roots after 24 hours of EST induction. (**G-L**) Imaging of $NSF_{WT}$:GFP (**G-I**) or $NSF_{EQ}$:GFP (**J-L**) and SynaptoRed™ C2 upon BFA application. (**M-R**) Effect on vacuolar traffic. Live imaging of $NSF_{WT}$:GFP (**M-O**) or $NSF_{EQ}$:GFP (**P-R**) and vacuolar soluble cargo marker AFVY:RFP in seedling roots.

Scale bars, 5 µm (**C**, **F**, **I**, **L**, **O**, **R**).

**Supplemental Table S1. Effects of dominant-negative NSF and αSNAP2 on seed viability**

| Parental lines (Female HZ x Male HM) | Viable seeds (%) | Non-viable seeds (%) | Total |
|---|---|---|---|
| UAS::$NSF_{WT}$:GFP #20-3 x RPS5A::GAL4 | 99 | 1 | 200 |
| UAS::$NSF_{WT}$:GFP #26-3 x RPS5A::GAL4 | 98 | 2 | 166 |
| UAS::$NSF_{EQ}$:GFP #2-2 x RPS5A::GAL4 | 48 | 52 | 125 |
| UAS::$NSF_{EQ}$:GFP #10-1 x RPS5A::GAL4 | 47 | 53 | 134 |
| UAS::RFP:$\alpha SNAP2_{WT}$ #1-2 x RPS5A::GAL4 | 99 | 1 | 198 |
| UAS::RFP:$\alpha SNAP2_{WT}$ #8-2 x RPS5A::GAL4 | 98 | 2 | 209 |
| UAS::RFP:$\alpha SNAP2_{LA}$ #2-1 x RPS5A::GAL4 | 45 | 55 | 164 |
| UAS::RFP:$\alpha SNAP2_{LA}$ #20-1 x RPS5A::GAL4 | 48 | 52 | 187 |

Heterozyous (HZ) maternal plants were crossed with homozyous (HM) paternal plants and the resultant F1 was analyzed for seed viability.

Expected viability values: 100% if dominant-negative NSF or αSNAP2 expression without deleterious effect, 50% if lethal.

**Supplemental Table S2. List of primers used for cloning**

| Primer name | Sequence (5'-3') | Construct |
|---|---|---|
| att1Bp-NSF-F | CAAAAAAGCAGGCTATGGCGGGTCGTTACGGA | 'pMDC7::NSF$_{\text{WT or E326Q}}$:GFP' |
| NSF_3xG_GFP_R | TCCTTTACTCATTCCTCCACCGCCAGTGAAGCGAATGAAGTC | 'pMDC7::NSF$_{\text{E326Q}}$:GFP' |
| NSF_3xG_GFP_F | CGCTTCACTGGCGGTGGAGGAATGAGTAAAGGAGAAGAACTT | 'pMDC7::NSF$_{\text{E326Q}}$:GFP' |
| GFP-stop_attB2p_R | ACAAGAAAGCTGGGTTCATTTGTATAGTTCATCCATGCC | 'pMDC7::NSF$_{\text{WT or E326Q}}$:GFP' |
| attB1p-mRFP-F | CAAAAAAGCAGGCTATGGCCTCCTCCGAGGACGTC | 'pMDC7::mRFP:$\alpha$SNAP2$_{\text{WT or L288A}}$' |
| mRFP_3xG_aSNAP2_R | ATGATCCCCCATTCCTCCACCGGCGCCGGTGGAGTGGCGGCC | 'pMDC7::mRFP:$\alpha$SNAP2$_{\text{WT or L288A}}$' |
| mRFP_3xG_aSNAP2_F | TCCACCGGCGCCGGTGGAGGAATGGGGGATCATCTGGTGAGA | 'pMDC7::mRFP:$\alpha$SNAP2$_{\text{WT or L288A}}$' |
| attB1 universal | GGGGACAAGTTTGTACAAAAAAGCAGGCT | 'pMDC7' (and sequencing) |
| attB2 universal | GGGGACCACTTTGTACAAGAAAGCTGGGT | 'pMDC7' (and sequencing) |
| NSF_Q326E_F | ATTTTTTGATGAGATTGATGCT | 'pMDC7::NSF$_{\text{WT}}$:GFP' |
| NSF_Q326E_R | AGCATCAATCTCATCAAAAAT | 'pMDC7::NSF$_{\text{WT}}$:GFP' |
| aSNAP2_A288L_attB2p_R | ACAAGAAAGCTGGGTCTATGTAAGGTCATCCTCCTC | 'pMDC7::mRFP:$\alpha$SNAP2$_{\text{WT}}$' |
| XhoI-NSF-F | AAAACTCGAGATGGCGGGTCGTTACGGA | 'pGIIB-UAS::NSF$_{\text{WT or E326Q}}$:GFP' |
| GFPstop-HindIII-R | TTTTAAGCTTTCATTTGTATAGTTCATC | 'pGIIB-UAS::NSF$_{\text{WT or E326Q}}$:GFP' |
| xhoI-mRFP-F | AAAACTCGAGATGGCCTCCTCCGAGGAC | ''pGIIB-UAS::mRFP:$\alpha$SNAP2$_{\text{WT or L288A}}$' |
| SNAP2wt stop-EV-R | TTTTGATATCCTATGTAAGGTCATCCTC | ''pGIIB-UAS::mRFP:$\alpha$SNAP2$_{\text{WT}}$' |
| SNAP2LA stop-EV-R | TTTTGATATCCTATGTAGCGTCATCCTC | ''pGIIB-UAS::mRFP:$\alpha$SNAP2$_{\text{L288A}}$' |
